# A surface metric and software toolbox for EEG electrode grids in the macaque

**DOI:** 10.1101/2020.03.10.986588

**Authors:** Fan Li, Tobias Teichert

## Abstract

**Background:** The past years have seen increased appreciation of non-invasive extracranial electroencephalographic (EEG) recordings in non-human primates (NHP) as a tool for translational research. In humans, the international 10-20 system or extensions thereof provide standardized electrode positions that enable easy comparison of data between subjects and laboratories. In the NHP, no such generally accepted, standardized placement system is available.

**New Method:** Here we introduce a surface metric and software package (NHP1020) that automates the planning of large, approximately evenly spaced electrode grids on the NHP skull.

**Results:** The system is based on one CT and one MRI image and requires the user to specify two intracranial markers. Based on this, the software defines electrode positions on the brain surface using a surface-based spherical metric similar to the one used by the international 10-20 system. The electrode positions are then projected to the surface of the skull. Standardized electrode grids can be shared, imported or defined with few high-level commands.

**Existing Methods:** NHP EEG electrodes are often placed on an individual basis relative to extracranial markers, or relative to underlying neural structures. Both approaches are time-consuming and require manual intervention. Furthermore, the use of extracranial markers in this species may be more problematic than in humans, because cranial muscles and ridges are larger and keep maturing long into adulthood thus potentially affecting electrode positions.

**Conclusion:** The key advantage of the current approach is the automated and objective identification of corresponding electrode positions in different animals. Automation was made possible by the use of a two-dimensional metric on the brain surface which has a simpler, i.e., more convex and sphere-like anatomy than the skull. This enables fast and efficient planning, optimization and calculation of large electrode grids.

## 1. Background

Recent years have seen a growing interest in non-invasive EEG-recordings in non-human primates (**NHP**) as a tool for translational research (e.g., Woodman et al., 2007; Sander et al., 2010; Godlove et al., 2011; Honing et al., 2012; Gil-da Costa et al., 2013; Purcell et al., 2013; Phillips et al., 2014; Rachalski et al., 2014; Gindrat et al., 2015; Itoh et al., 2015; Teichert, 2016; Teichert et al., 2016; Teichert, 2017; Holliday et al., 2017; Teichert et al., 2019; Itoh et al., 2019). Despite the availability of intra-cranial measures of neural activity that are more spatially-specific, NHP EEG has a number of appealing properties. (1) It has the ability to serve as a bridge method that can be used in both humans and NHPs, thus increasing the translatability of invasive intracranial recordings that can only be performed in NHPs (Arthur & Starr, 1984; Paller et al., 1988, 1992; Sander et al., 2010; Purcell et al., 2013; Teichert et al., 2019). (2) In some cases, a more global measure of brain activity that sums over activity in many different brain regions may actually be more useful than highly localized recordings– for example, when determining either sleep state (Rachalski et al., 2014), or distributed signals such as mismatch negativity or the P3 (Gil-da Costa et al., 2013; Teichert et al., 2019). (3) The use of sophisticated analysis tools in combination with detailed anatomical models of the head can provide accurate source localization to track neural reorganization over time (Gindrat et al., 2015), or to guide subsequent invasive intracranial recordings or microinjections.

In order to take full advantage of the translational potential of this technique, it is essential to optimize the recording platform used for NHP EEG. Techniques in humans have been refined over many decades, but NHP EEG is still in its relative infancy. In humans, EEG electrodes are typically placed according to the international 10-20 system (Jasper, 1958) or extensions thereof (Oostenveld & Praamstra, 2001; Chatrian et al., 1985). The 10-20 system specifies electrode location based on four extracranial landmarks (inion, nasion and the two pre-auricular points) and measurements between these points (Jasper, 1958). The generally accepted, standardized use of the 10-20 system assures comparable and reliable electrode placement across subjects and laboratories. It also assures approximately even coverage across accessible parts of the head, and can easily be expanded to yield either higher coverage density, or to cover parts of the anatomy that were previously not used for EEG recordings (Oostenveld & Praamstra, 2001; Chatrian et al., 1985). The use of EEG caps have eliminated the need for time-consuming measurements to identify the correct electrode locations. The electrode positions of the 10-20 system can conveniently be specified in a two-dimensional spherical coordinate system in which Cz serves as the north pole.

In the NHP, no such generally accepted, standardized system is available. Some labs have used extracranial homologs of the Inion, Nasion, and the pre-auricular points to identify electrode position using extracranial measurements, very much like in humans before the advent of electrode caps (Woodman et al., 2007; Honing et al., 2012; Itoh et al., 2015). In other instances, electrodes are placed on an individual basis relative to neural structures such as a particular sulcus or gyrus. This individual electrode approach is particularly useful when placing sensors such as sub- or epi-dural ECoG electrodes that sample activity from a spatially restricted region of the brain (Freeman et al., 2003). However, the activity of extracranial EEG electrodes depends on neural activity in the entire brain. Hence, it is likely less meaningful to determine their position relative to a single gyrus that may show substantial variability relative to other structures in the brain. Rather, it may be more meaningful to determine electrode position relative to overall gross brain anatomy. A second limitation of the individual electrode approach is that it is difficult to automate, and hence takes a significant amount of time when electrode counts exceed certain limits.

To date, there is no existing, generally accepted surface-based 2-dimensional coordinate system that enable the sharing of electrode positions between labs or the automated generation of large, approximately evenly spaced electrode grids on the NHP skull. A surface-based metric has been difficult to implement, because the skulls of NHPs are not as smooth and convex as those of humans. Most notably, the protruding orbital bone and sharp occipital ridge can affect the extracranial measurements that underlie the human 10-20 system. We have circumvented the complex cranial anatomy by using intracranial markers and measurements to find electrode positions on the brain surface before projecting them to the closest point on the skull. This two-dimensional metric allows us to identify identical electrode positions on different animals and easily define entire grids of approximately evenly spaced electrodes. The proposed brain-based surface-metric also allows us to import entire electrode grids, such as the international 10-20 system, that are specified in spherical coordinates. Similar to the metric underlying the 10-20 system it can easily be defined based on a limited number of anatomical landmarks that can reliably be identified in any animal.

This manuscript outlines the technical principles of the method in more detail, and introduces a Matlab-based toolbox (**NHP1020**) that allows implementation of these ideas. It then showcases a couple of electrode grids generated with the software, as well a auditory evoked potentials recorded from one specific electrode grid that was implanted based on the electrode positions identified with the software.

## 2. Methods

### 2.1 Requirements

The NHP1020 software as presented here requires two types of images from each animal: a T1 weighted MR image and a CT image (**Fig 1**). The former is used to identify two neural landmarks, the later is used to extract a surface model of the skull. Both images need to be aligned in a stereotaxic coordinate frame, i.,e., with the origin corresponding to the inter-aural point and the z-plane defined by the center of the two ear-canals and the lowest point of the left (or right) eye-socket. It may be possible to adapt the software to use either the MR or the CT image only. However, having both images facilitates the process.

**Figure 1:**
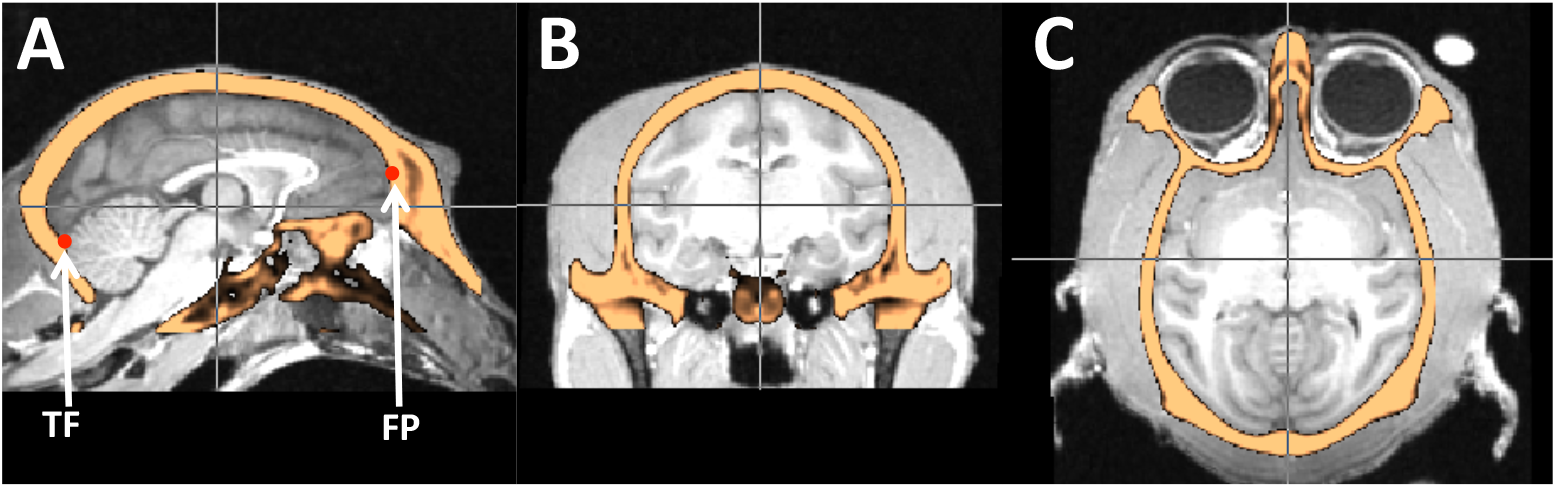
Requirements of the NHP1020 software. The NHP1020 toolbox assumes that users have one MR T1 (greyscale) and one CT image (orange) of the animal in question aligned to a stereotaxic coordinate frame. In addition, the user needs to extract a mask of the brain and manually define two landmarks on the midline sagittal plane: the caudal end of the transverse fissure (TF) and the frontal pole (FP) as indicated by the red dots.

### 2.2 Landmarks

The NHP1020 software calculates electrode locations based on two landmarks. The first landmark is the caudal end of the transverse cerebral fissure (**TF**) on the midline (**Fig 1A**). The second landmark is the frontal pole of the cerebrum (**FP**), also on the midline. These two points play similar roles as inion and nasion of the international 10-20 system. Note that while these points serve the same functional role in setting up the electrode positioning system, they do not refer to homolog parts of the anatomies. Both landmarks need to be identified by the user in the stereotactically aligned images and passed to the software.

### 2.3 Preprocessing of MR image

The brain extraction routine (**bet**) from **FSL** (Woolrich et al., 2009; Smith et al., 2004; Jenkinson et al., 2012) was used to generate a mask of the brain-volume from the T1 image. Masking the T1 images with the aligned and suitably thresholded CT image can increase the success rate of the **bet** routine to come up with the correct segmentation of the T1 image. If needed, the brain mask image can be edited by hand. Alternative approaches can be used to generate a mask of the brain volume. For example, it may be possible to use the CT image to create an endo-cast of the skull cavity, which should yield virtually identical electrode placements.

### 2.4 Surface models

As a first step, the NHP1020 software reads in the MR-defined brain-mask image and the raw CT image of the skull. Using suitable user-defined threshold values, the program uses the isosurface function to generate a surface model of the brain and the skull. In addition, the program uses the reducepatch function to reduce the number of patches in the surface models to a more manageable number.

### 2.5 Surface-based coordinate frame

The NHP1020 software defines two one-dimensional manifolds on the brain surface model (**Fig 2A**). The first manifold corresponds to a cut of the brain volume along the sagital plane, ie, ml=0. We refer to this manifold as the midline. We then identify the two points on the midline that correspond to the user-defined coordinates of TF and FP. The software then measures the length of the midline manifold between TF and FP. It then defines homologs of electrode positions Oz and Fpz by moving 10% and 95%, respectively, of the distance from the TF towards FP. In our coordinate frame further defined below, these two electrodes define the positions (0,0) and (1,0), respectively. In this context we refer to electrodes by these coordinates, to avoid confusion with naming convention of the international 10-20 system.

**Figure 2:**
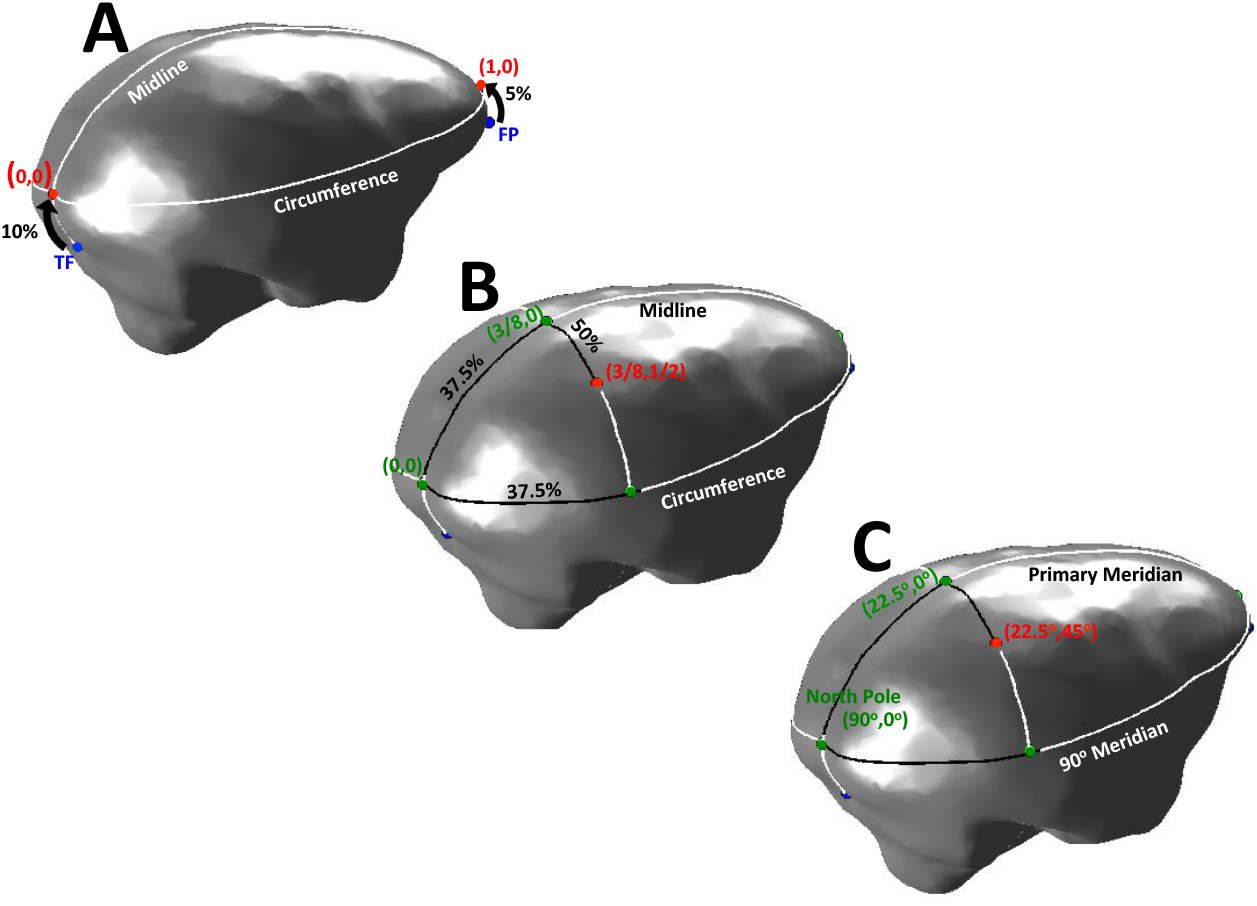
Construction of the NHP surface-based coordinate frame. (**A**) The NHP1020 system is based on two landmarks: the caudal end of the transverse cerebral fissure (**TF**) and the frontal pole of the cerebrum (**FP**). On the midline, we define the points (0,0) and (1,0), as the 10% and 95% mark between TF and FP. (**B**) The position of any electrode on the brain surface can be defined in the following 2-dimensional surface-based coordinate frame. The first value defines the anterior-posterior position of the electrode as a fractional value. In the example, the fraction is 3/8=37.5%. This value defines three points: one point on the midline, 37.5% the distance from (0,0) to (1,0), and two points on the circumference, 37.5% from (0,0) to (1,0) clockwise as well as counterclockwise. These three points define a plane that intersect the brain surface to define a one-dimensional manifold. The second coordinate of the electrode determines its position on this manifold. A value of 0 corresponds to the midline, values of +1 and −1 correspond to the intersections of the manifold with the circumference. The example shows an electrode halfway between the midline and the intersection with the equator over the right hemisphere with the coordinates (3/8,1/2). See section 2.5 for details.

The second manifold arises by cutting the brain-mask surface model along a plane with zero roll that crosses through (0,0) and (1,0). In other words, the plane is defined such that the medio-lateral coordinate of its normal vector is zero. We refer to the cut along this plane as the circumference (**Fig 2A**). The circumference corresponds to the line in the international 10-20 system that loops around the skull from Fpz via T7, Oz, T8 and back to Fpz.

We can now set up additional support manifolds to mediate electrode placements. The additional support manifolds are defined by cuts of the brain surface-model with specific planes. Each plane is defined by a single value that determines its anterior-posterior position (ap-value). The scalar ap-value ranges between 0 and 1 and defines three points on the surface model. These three points in turn define the plane. For example, the support manifold with ap-value 0.5 defines the points half way between (0,0) and (1,0) on the midline (which in the human 10-20 system would correspond to Cz) and the points half way between (0,0) and (1,0) on the circumference going clockwise (1/2,1) (which would correspond to T7) and counterclockwise (1/2, −1) (T8) (**Fig 2B**).

These support-manifolds define a two-dimensional coordinate system in which to specify the position of any electrode on the brain surface (**Fig 2B**). The first coordinate defines the anterior-posterior position of the support manifold (ap-value). The second coordinate defines the medio-lateral electrode position on this support manifold (ml-value). The coordinates (3/8,0) index a position on midline the three eighths of the way from (0,0) to (1,0) (**Fig 2B**). The coordinates (3/8,1/2) correspond to a position on the same support manifold, but half way down over the right between (3/8,0) and (3/8,1). The coordinates (3/8, −1/2) would correspond to the same position over the left hemisphere.

### 2.6 Translation into spherical coordinates

It important to note that this two-dimensional coordinate system is very closely related to a spherical coordinate frame with Oz and Fz as North and South pole, and the midline as the primary meridian (**Fig 2C**). All points on the dorsal half of the brain, i.e., all points that lie above the circumference are uniquely defined by their elevation and their azimuth. The example point (3/8,1/2) corresponds to an elevation of 22.5 degrees and an azimuth of 45 degrees. In general, the point (x,y) with 0 ≤ *x* ≤ 1 and −1 ≤ −*y* ≤ 1, corresponds to an elevation of 180*(0.5−x) degrees and an azimuth of 90*y degrees.

This translation into a spherical coordinate frame is important because electrode positions in humans are often specified in spherical coordinates. However, in humans the spherical coordinate frame is set up such that the north pole is located at Cz, not Oz as defined here for the non-human primates. Nevertheless, human electrode positions, such as the ones of the 10-20 system, can easily be imported and projected onto the surface of the non-human primate brain using the NHP1020 software.

The deviation from a true spherical coordinate frame are minor and mostly evident for points close to the +/− 180 degree meridian, i.e., close to the midline on the ventral side of the brain. Since such electrode positions are highly unusual in any positioning scheme, these deviations are of no concern for common electrode positions in the 10-20, 10-10 or 10-5 system.

### 2.7 Specifying electrode grids

A key feature of the NHP1020 software is its ability to easily specify large electrode grids that contain approximately equi-distantly spaced electrodes. This is accomplished in one of two ways. Pre-specified layouts such as the international 1020 system can be imported from eeglab (Delorme & Makeig, 2004) location files. **Figure 3A** shows the international 1020 system projected onto an example non-human primate brain mask.

**Figure 3:**
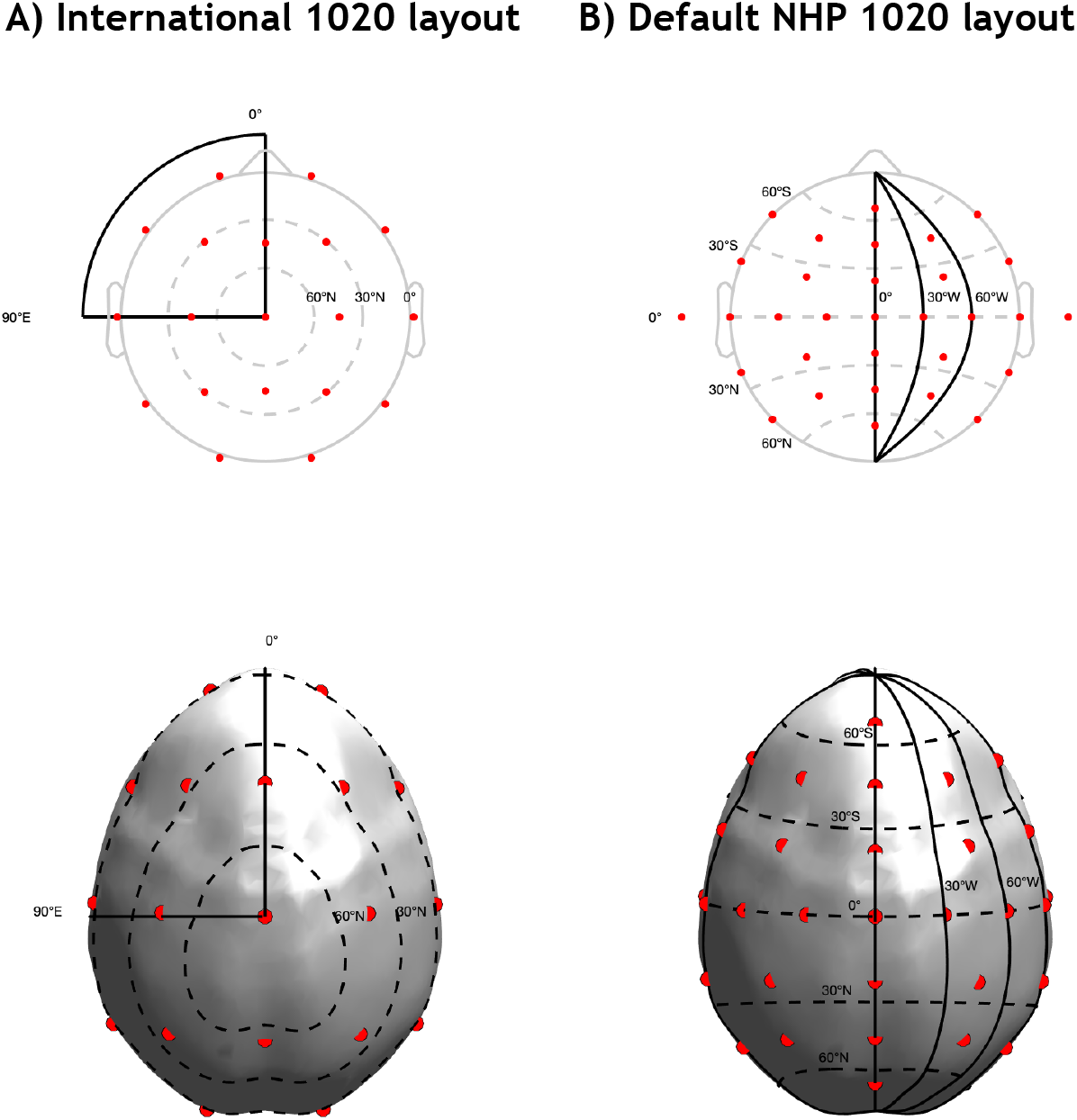
Comparison of two electrode layouts. (**A**) International 10-20 system projected on the monkey brain using the NHP1020 spherical coordinate system. (**B**) The default NHP1020 layout.

Alternatively, grids can be specified using a high-level short-hand description that can be provided to the software in a text file. The short-hand consists of several lines of text each containing of two positive integers *n* and *m* separated by a comma. The number of lines corresponds to the number of support manifolds on which to create electrodes. For example, in a file with 9 lines, the first line describes the most posterior, the last one the most anterior manifold.The ap positions of the manifolds are equally spaced at positions at ap positions of 0/8, 1/8, 2/8,…, 8/8. The values *n* and *m* define the number and spacing of electrodes on that manifold. For example, the line ‘`n,m`’ corresponds to a pattern in which the electrodes are located on the positions E(*x*, *j/m*) with −*m* ≤ *j* ≤ *m*.

**Figure 3B** shows the default 31 channel coverage pattern that is used by the NHP1020 software. It uses 9 evenly spaced support manifolds, i.e., ap=0/8, 1/8, … 8/8. The first and last manifolds are placeholders and have no electrodes on them (‘`0,0`’). The second and second to last manifolds have one electrode each positioned right on the midline (‘`0,1`’). The third, forth, sixth and seventh manifolds have five electrodes each (‘`2,2`’). The fifth manifold has nine electrodes (‘`4,3`’). Overall, this grid describes 31 electrode positions. One of these electrodes is typically used as reference, leaving an additional 2 electrodes when using standard 32 channel recording amplifiers. The remaining two electrodes can be positioned on the orbital ridge to facilitate the detection and removal of blinks and eye-movement artifacts.

### 2.8 Projection to the skull surface

In the last step, the software projects the electrode positions that were defined on the brain to the skull. To that aim, the software computes a surface model of the skull and then determines for each electrode position on the brain the closest position on the skull. Alternatively, the software can determine the normal vector of the brain surface at the electrode position and then identify the intersection of the normal vector with the skull. By default, the NHP1020 software implements the first method because it provides more robust results for some electrode positions.

### 2.9 Working with the NHP1020 GUI

The NHP1020 software allows the user to design electrodes grids for NHPs following the methodology outlined above. After navigating matlab to the ‘NHP1020GUI’ folder, the main GUI can be opened with the command ‘NHP1020’. The workflow is structured into four main steps which are reflected in the four main panels of the GUI (**Figure 4)**.

**Figure 4:**
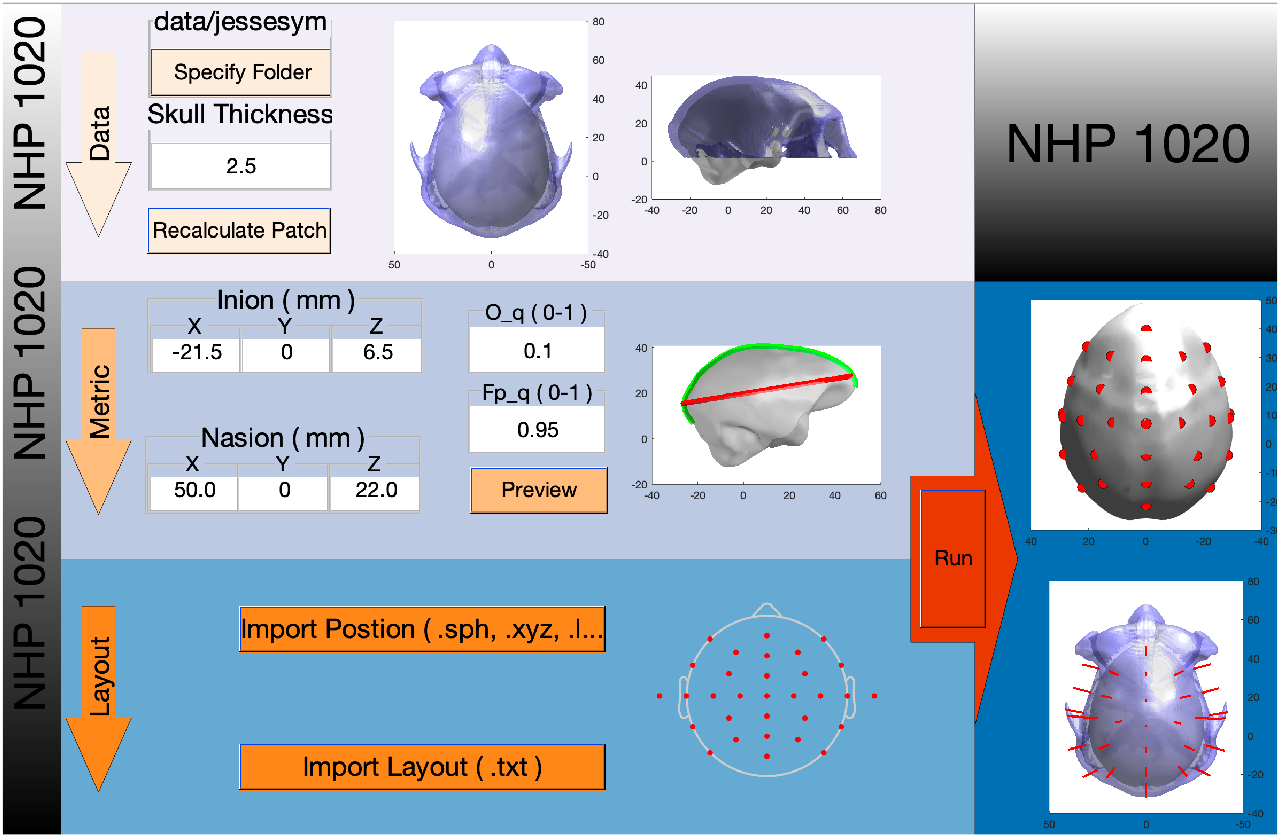
NHP1020 GUI. The ‘**Data**’ panel consists of two buttons. The ‘Specify Folder’ button allows users to specify the main data and results folder. The ‘Recalculate Patch’ button prompts the software to recalculate and display the brain and skull surface patches. Once generated the surfaces are displayed in two figures (Left: brain and skull top view, Right: brain and skull side view). The ‘**Metric**’ panel requires users to specify the position of Inion and Nasion. While generally discouraged, it also provides the option to change O q or Fp q. The preview button prompts the software to calculate and display midline and circumference in a sagital view of the brain surface. The ‘**Layout**’ panel allows users specify the desired electrod locations.The ‘Import Position’ button imports electrode positions specified in any format that can be read by eeglab (’.sph’,’.xyz’,’.loc’). The ‘Import Layout’ button opens text files that contain electrode locations specified in the NHP1020 shorthand. Once specified, the electrode positions are displayed in cartoon format using the eeglab display function. The ‘**Run**’: After loading the anatomical files, specifying the metric and the desired electrode layout, the user can press the ‘Run’ button to calculate the electrode positions on the brain (top panel) and project them to the skull (bottom panel).

#### (A) Data

The main purpose of the data panel is to specify the data folder that contains the CT image and the brain mask in nifty format. This folder will be used to store all the intermittent data and the final results of the process. The software distribution of NHP1020 contains a fully functional example data set that is located in the folder `NHP1020/data/jessesym`. Once a folder is specified, the GUI will look for two files that contain the character sequence ‘*CT_skull*.nii.gz’ and ‘*inskull*.nii.gz’. If two nifty files match that description, the software will open them and create surface patches of the skull and the brain and save them in stl format. If the necessary stl files have already been generated in an earlier session, the software will immediately open the stl files, rather than generating them from scratch because this process is time-consuming. As soon as the patches have been generated or loaded, they will be automatically displayed in the figures to the right of the data tab. The skull surface is plotted in light blue. The gray brain surface is visible through the semi-transparent skull. For some animals, the default settings may result in holes in the skull. This can typically be resolved by lowering the parameter ‘Skull Thickness’ and pressing the ‘Recalculate Patch’ button. The steps of the data tab can be considered complete once intact skull and brain surfaces have been generated and displayed.

#### (B) Metric

The main purpose of the ‘metric’ tab is to set up the two-dimensional surface coordinate frame that lies at the heart of the NHP1020. To do so, the user needs to specify the positions of NHP’s Inion- and Nasion-homolog as defined in **section 2.5**. We suggest identifying these points in a dedicated nifty file viewer such as fsl_eyes. The GUI also provides the option to change the values of O_q and Fp_q. However, it is suggested to use the default values for consistency across labs. Once specified, the ‘Preview’ button will display a figure of the brain patch with the current midline and circumference superposed. If the metric is acceptable, the user can proceed to the layout tab.

#### (C) Layout

The layout tab allows users to specify the electrode grid they want to project onto the monkey skull. Layouts can be imported from eeglab location files in .sph, .xyz. and .loc formats. A couple of sample grids such as the 10-20 and 10-10 system in spherical coordinates are included in the ‘template’ folder of the NHP1020 software. Alternatively, grids can be specified using the NHP1020 shorthand described in section 2.8. A text file containing the shorthand definition of the default NHP1020 layout is also provided in the ‘template’ folder. Once a layout has been loaded, the software displays a flat-map mock-up of the layout.

#### (D) Run

Once the layout has been finalized, the electrode positions can be projected to the rhesus skull by pressing the ‘Run’ button. This process is time-intense, so it makes sense to carefully choose the layout before identifying electrode positions on the individual monkey brain and projecting them to skull. The results are saved in a text file (’positionList ISI.txt’) in the main folder that was specified in the data tab. Each row in the file represents an electrodes by features (Number: row number, Name: electrode’s label, brainX: electrode’s x-axis value on the brain, brainY: electrode’s y-axis value on the brain, brainZ: electrode’s z-axis value on the brain, skullX: electrode’s x-axis value on the skull, skullY: electrode’s y-axis value on the skull, skullZ: electrode’s z-axis value on the skull). A second, more detailed result file also specifies the normal vectors at each of the electrode positions on the skull. This information can be used to identify the settings of a stereotaxic device that allows a normal approach to the skull. This is particularly helpful when positioning electrodes close to the circumference. An add-on piece of software that translates the electrode positions and their normal vectors into settings of the Kopf micro-manipulator is included in the ‘r-files’ folder of the distribution of NHP1020.

## 3. Results

We have currently implanted electrode grids that were planned with the NHP1020 software in four rhesus macaques. The treatment of the monkeys was in accordance with the guidelines set by the U.S. Department of Health and Human Services (National Institutes of Health) for the care and use of laboratory animals. All methods were approved by the Institutional Animal Care and Use Committee at the University of Pittsburgh.

The electrodes were manufactured either from medical grade titanium or medical grade stainless steel. The diameter of the electrodes was 3 mm to increase the contact surface between skull and electrode (**Fig5A**). The sides of the electrodes contained grooves to increase stability after oseo-integration. The top side of each electrode contained a 1mm diameter 2mm deep hole. Inside this hole we inserted a 1mm diameter amphenol pin. The excess part of the pin was cut off and covered with a blob of solder. Teflon-coated stainless steel wire was soldered to the amphenol pin and routed to a 36 channel omnetics connector. Electrodes were implanted in non-penetrating 1 mm deep flat-bottom holes that were drilled with the help of a flat-bottom carbide end-mill (**Fig 5B**). After installation, most of the wires and electrodes were covered in dental acrylic (**Fig 5C**). Only some of the electrodes close to and below the circumference remained outside of the acrylic head cap (**Fig 6)**. The omnetics connector was partially embedded in the acrylic and positioned inside a small PEEK connector box that could be covered with a lid to prevent dirt from getting into the connector.

**Figure 5:**
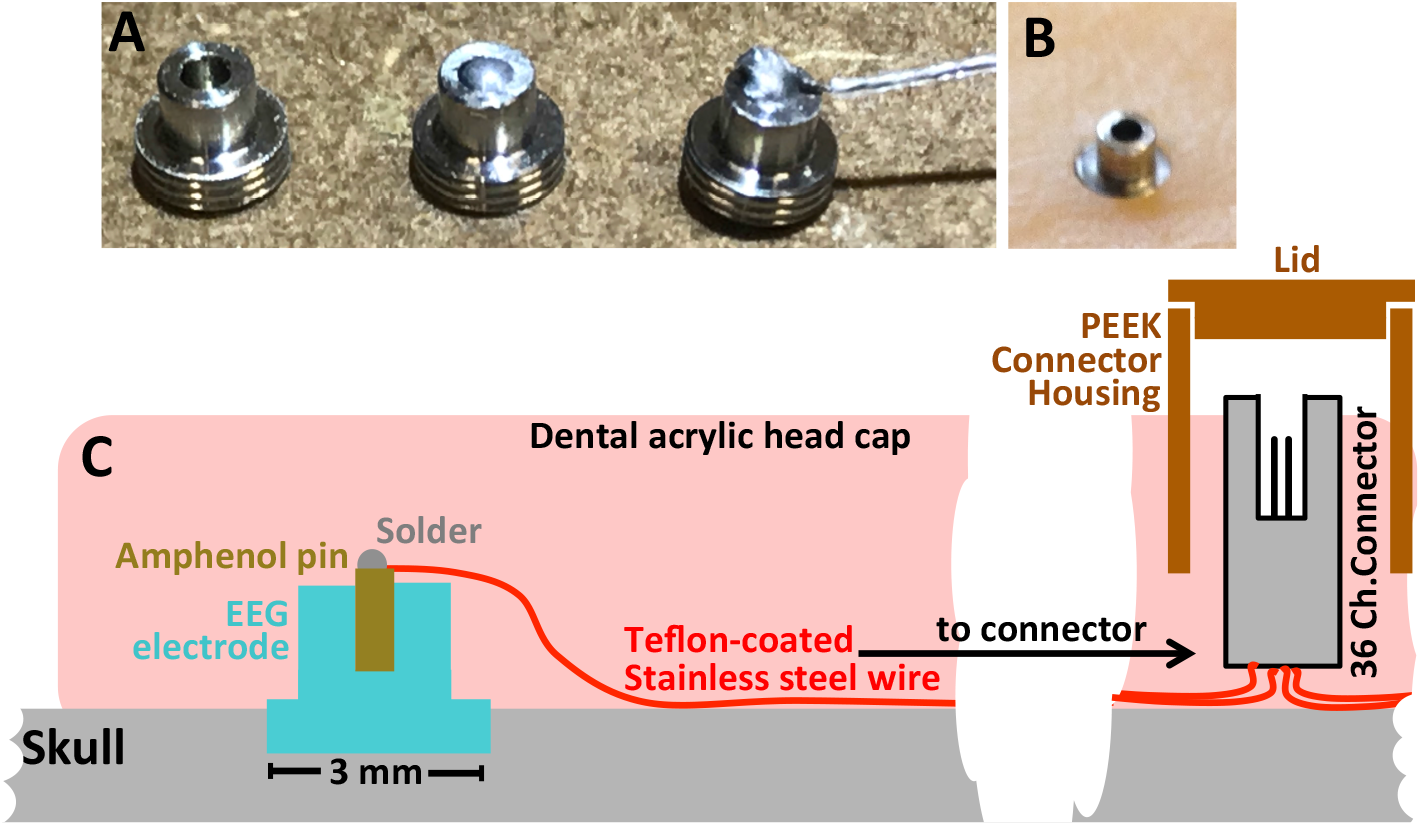
Implantation. (**A**) Left: in-house manufactured electrode. Middle: after insertion of amphenol pin and solder drop. Right: after attachment of stainless steel wire. (**B**) Electrode embedded in flat-bottom hole. (**C**) Cartoon of electrode implantation and wiring scheme.

**Figure 6:**
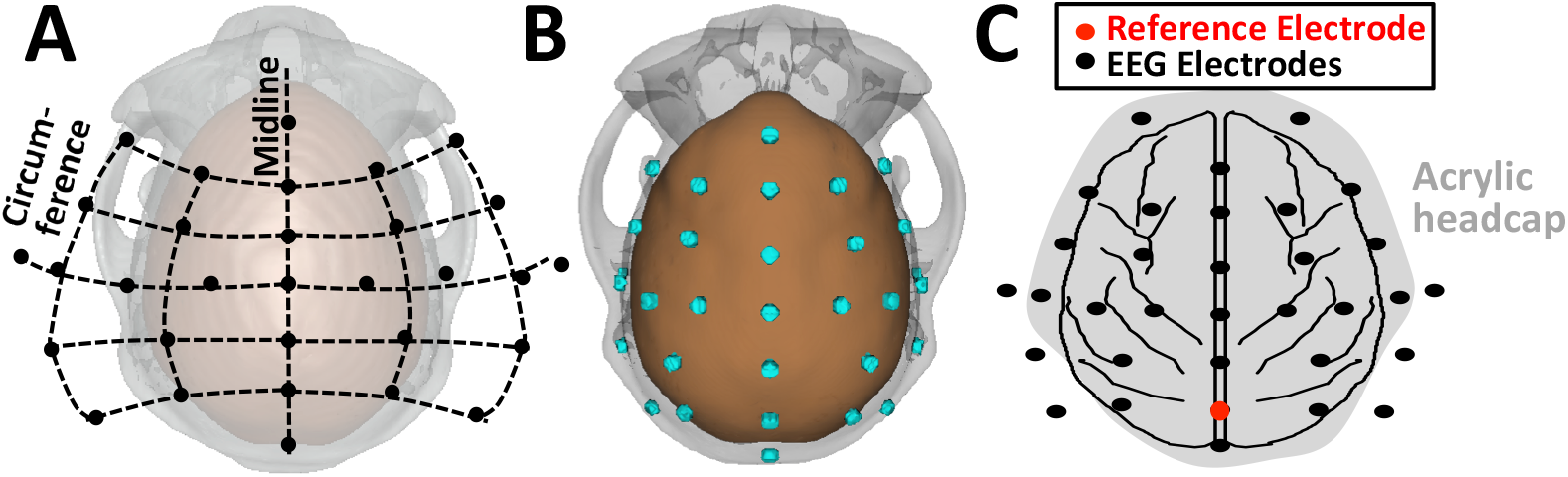
32 channel example coverage. (**A**) The default NHP1020 electrode grid in flat-map format. (**B**) Electrode grid viewed from the top. (**C**) Position of 33 implanted electrodes including one reference electrode in one example animal.

**Figure 6** shows an overview of the electrode grid in one animal that used the default NHP1020 grid outlined above. The two additional electrodes were positioned over the orbital bone to monitor eye-movements and blinks. Electrode E(2/8,0) served as the reference electrode during recording. Offline, the signals were re-referenced to E(1/8,0)

**Figure 7** shows auditory evoked potentials recorded from this electrode array in response to 50 ms long sinusoidal tones. A total of 8 EEG components could be defined based on different timing and spatial distribution over the head (Teichert, 2016). It is noteworthy that the evoked activity shows strong similarity for adjacent electrodes, a finding that is consistent in all animals. This is in contrast to intracranial recordings with epi-dural recordings where patterns of activity reflect more local processing and can vary with a higher spatial frequency (Freeman et al., 2003). In this animal, evoked potentials tended to be larger over the left hemisphere. A similar asymmetry was not found in three other animals. With the exception of overall amplitude differences, electrodes on corresponding part of the two hemispheres show highly similar time-courses, more so than adjacent electrodes on the same hemisphere. More in-depth analyses of these EEG data have been presented in earlier work (Teichert et al., 2016; Holliday et al., 2017; Teichert, 2017; Teichert et al., 2019).

**Figure 7:**
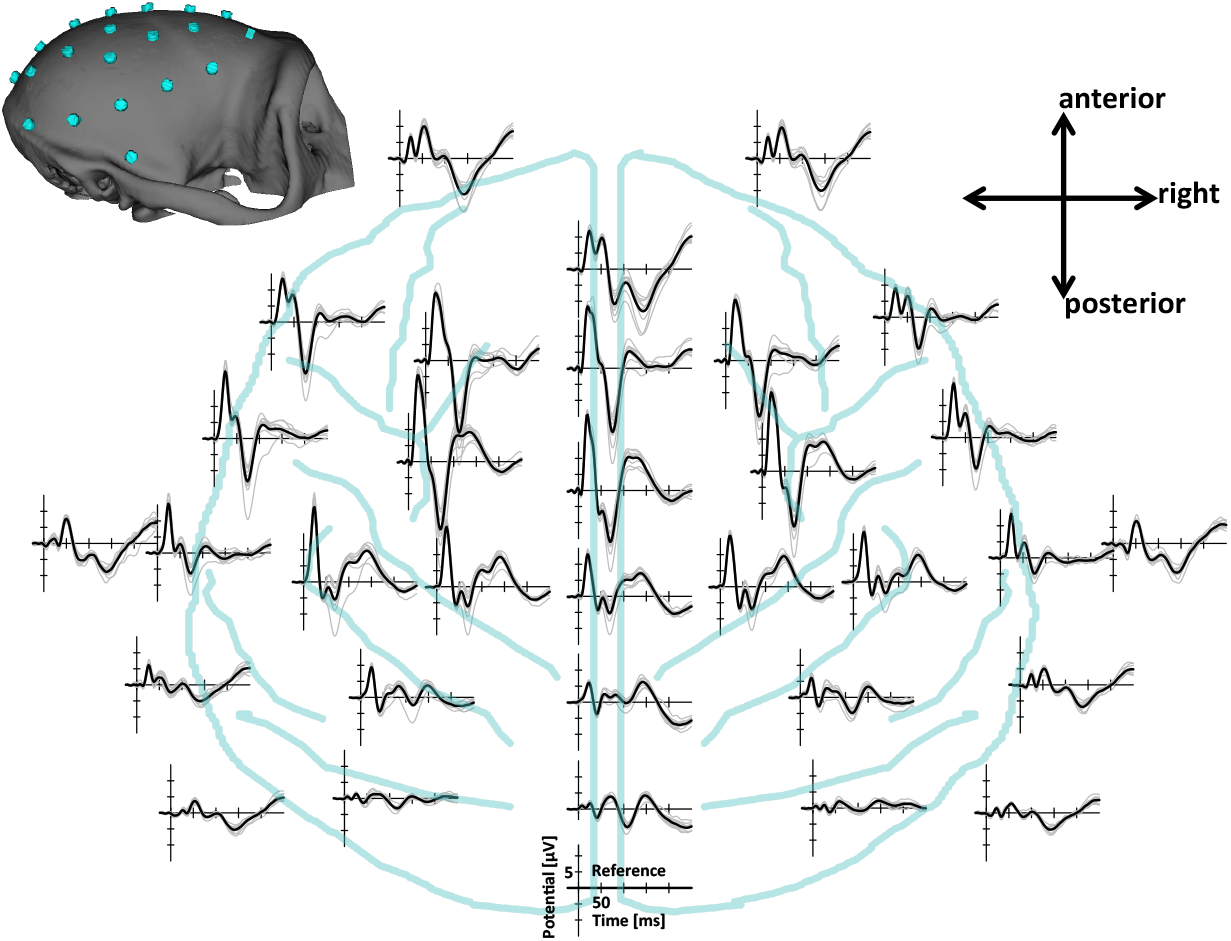
Auditory evoked responses. Sample traces from 32 electrodes in response to the onset of pure tones at time zero. The underlying light blue traces indicate the brain outlines and approximate location of the main sulci in the rhesus viewed from above. Thick black lines indicate the mean over 15 recording sessions. Thin lines indicate the mean of individual recording sessions.

## 4. Discussion

The past years have seen increased appreciation of non-invasive extracranial EEG recordings in non-human primates as a tool for translational research. To facilitate EEG recordings in NHPs, we proposed a unifying surface-based metric and an automated algorithm that allows users to plan large EEG electrode grids that can be projected to equivalent locations on different animals. The software has four main features. 1) Electrode locations can be calculated automatically without manual intervention. 2) Even large electrode grids can be planned, designed and reviewed within minutes. 3) It defines a unifying placement scheme that allows comparable electrode positions between animals and labs. 4) It can import electrode positions such as the international 1020 system in spherical coordinates and project them onto the monkey skull.

Conceptually the NHP1020 provides an interesting alternative to positioning schemes based on local anatomy. As extracranial EEG electrodes are affected by many dipoles distributed all over the brain, a positioning scheme that only takes the anatomy of the immediate surrounding into account (e.g, “electrode was placed above FEF”) may render less comparable signals between animals than a system that takes more of the brain anatomy into account. That said, both schemes of placing electrodes and/or describing electrode position have their own advantages. The most appropriate approach needs to be determined on a case-by-case basis to meet the demands of each specific application.

### Alternatives to NHP1020

Several labs have used extracranial homologs of the Inion, Nasion, and the pre-auricular points to identify electrode position using extracranial measurements (Woodman et al., 2007; Honing et al., 2012; Itoh et al., 2015). The key appeal of this approach is the direct homology to the 10-20 electrode placement schemes used in humans. However, to our knowledge this approach has not be automated and is thus more time-consuming. We had decided to base our automated algorithm on intracranial measurements for several reasons: 1) They are simpler because the intracranial anatomy is more convex and sphere-like; 2) They are more proximal because electrode positions are independent of skull ridges and muscles which are substantially larger and continue to mature for longer in monkeys compared to humans. Because CT and MRI images are so commonplace that it is not necessary to use more indirect and distal extracranial markers. That said, we believe that both approaches, extracranial and intracranial, should provide unique electrode positions and thus a mapping between the two. Thus, neither of the two methods will provide ‘better’ electrode positions. The choice between the two approach will most likely be driven by individual preference, convenience and availability of MRI/CT images, as well as whether electrodes will be mounted on the scalp or the skull.

The company Easy Cap has recently developed an EEG cap with 32 electrodes for macaque monkeys (Gindrat et al., 2015). Their cap uses concentric electrode positions around Cz. While this is a great option for NHP scalp EEG it is less useful to determine EEG electrode location for implantation in the skull.

### Known limitations

The main limitation of the NHP1020 approach is that there is no strong theoretical rational for the choice of landmarks and the definition of the positions (0,0) and (1,0). Other landmarks could be used to define a two-dimensional surface-based coordinate system. The only way one particular coordinate frame could prove superior to others is if it leads to more consistent EEG signals for electrodes placed at identical positions in different animals. Given the very low number of subjects typically used in NHP experiments and the high inter-subject variability in brain anatomy and function, meaningful empirical data that speaks to this question is unlikely to be available soon.

### Disclaimer

The NHP1020 software provides a method to identify similar positions on different rhesus monkeys. It uses a similar approach as the international 10-20 system and variants thereof, but it will not necessarily identify electrode positions that are homolog to the human, neither with respect to their position relative to neural structures or generators, nor with respect to the signals recorded from these electrodes. If they exist, such functional homologies will need to be established either empirically or via simulations using detailed brain anatomy and realistic head-models.

## Code

The most up-to-date version of the NHP1020 software is available for download at https://github.com/Teichert-Lab/NHP1020. Use the following command to download: git clone https://github.com/Teichert-Lab/NHP1020 NHP1020

## Notes

https://github.com/Teichert-Lab/NHP1020

